# Dopaminergic Changes in the Subgenual Cingulate Cortex in Dementia with Lewy Bodies Associates with Presence of Depression

**DOI:** 10.1101/2024.01.09.574871

**Authors:** Lina Gliaudelytė, Steven P Rushton, Rolando Berlinguer-Palmini, Alan J Thomas, Christopher M Morris

## Abstract

In addition to the core clinical features of fluctuating cognition, visual hallucinations, and parkinsonism, individuals with dementia with Lewy bodies (DLB) frequently experience chronic and debilitating major depression. Treatment of depression in DLB is hampered by a lack of available effective therapies and standard serotonergic medication for major depressive disorder (MDD) is typically ineffective. Dysfunction of dopaminergic neurotransmission contributing to anhedonia and loss of motivation has been described in MDD. The subgenual anterior cingulate (sgACC) is important in mood regulation and in the symptomatic expression of depression, displaying structural, functional and metabolic abnormalities in MDD. To assess dopaminergic and serotonergic synaptic changes in DLB, post mortem sgACC tissue from DLB donors with and without depression was investigated using high-resolution stimulated emission depletion (STED) microscopy, as well as Western and dot blotting techniques. STED imaging demonstrated the presence of α-synuclein within individual dopaminergic terminals in the sgACC, α-synuclein presence showing a significant positive correlation with increased SNAP25 volumes in depressed DLB cases. A reduction in dopaminergic innervation in the sgACC was observed in DLB cases with depression, along with reduced levels of multiple dopaminergic markers and receptors. Limited alterations were observed in serotonergic markers. Our work demonstrates a role for dopaminergic neurotransmission in the aetiology of depression in DLB. Careful and selective targeting of dopaminergic systems may be a therapeutic option for treatment of depression in DLB.

## Introduction

Dementia with Lewy bodies (DLB) is a significant cause of morbidity in older populations, representing between 5-15% of all clinical dementia cases with some clinicopathological studies showing over 20% (McKeith et al. 2005; McAleese et al. 2020). Clinically, DLB is characterised by core symptoms of fluctuating cognition, parkinsonism, the presence of recurrent complex visual hallucinations and REM sleep behaviour disorder (McKeith et al. 2017). Many individuals with DLB also show additional psychiatric features at high prevalence, with co-morbid major depression a feature of DLB in 50-80% of cases (Stefanova et al. 2000; Yamane, Sakai, and Maeda 2011; Jellinger 2023; Kuring, Mathias, and Ward 2018). Depression is also often present at prodromal DLB stages and persists throughout the clinical course (Fujishiro et al. 2015; Kuring, Mathias, and Ward 2018). Depression in dementia is associated with poor quality of life, increased morbidity and more rapid cognitive decline and consequently treatment of depression would have significant patient benefit (Lenze et al. 2005; Starkstein et al. 1992; Breitve et al. 2016; Boot et al. 2013; Gatt et al. 2017; Terum et al. 2021; Jiang et al. 2023; Christensen, Schmidt, and Grande 2023; Joshi et al. 2022).

The subgenual anterior cingulate cortex (sgACC) is a key brain region associated with mood and anxiety disorders, which relays emotional information between limbic, cortical and subcortical regions (Greicius et al. 2007; Scharnowski et al. 2020). Abnormalities in sgACC activity, connectivity and grey matter volume have been shown in depressive disorders (Drevets, Savitz, and Trimble 2008; Wu et al. 2016; Drevets 2001). The sgACC has been used as a target region for treating treatment resistant depression with deep brain stimulation (Berlim et al. 2014; Rizvi et al. 2011; Holtzheimer et al. 2012). Both the sgACC and amygdala are highly inter-connected and are thought to be a part of an automatic emotion regulation circuit (Johansen-Berg et al. 2008; Davey et al. 2015).

The monoamine hypothesis of depression is well established in unipolar major depressive disorder (MDD), suggesting that reduction primarily in serotonergic and noradrenergic neurotransmission underpins depressive symptoms (Hirschfeld 2000), with most antidepressants modulating serotonin and noradrenaline activity. There is considerable evidence in support of the monoamine hypothesis in individuals with MDD, with reduced 5HT concentrations observed in serum and reduced 5HIAA in CSF in patients with MDD, along with decreased serotonin 5HT1_A_ and 5HT2_A_ binding in anterior cingulate cortex (ACC) in MDD (Wang et al. 2016; Baeken, De Raedt, and Bossuyt 2012). Serotonergic neurotransmission may be impaired in DLB compared to AD patients with depression, with reduced 5HT and metabolite concentrations observed in limbic regions (Vermeiren et al. 2015). However, reductions in levels of 5HT as a basis of depression have been challenged by several studies, primarily based on reduced and delayed response to 5HT based therapies (Lacasse and Leo 2005; Andrews et al. 2015; Ruhe, Mason, and Schene 2007; Karg et al. 2011). Data on efficacy of serotonergic treatment of depression in DLB is lacking, and in depression in Parkinson’s disease (PD), serotonergic treatment shows minimal effects in a few small-scale trials (Torun, Bayulkem, and Torun 2010; Barone et al. 2006).

Dopamine plays a role both in reward and stress (Pignatelli and Bonci 2015) with dysfunction of dopaminergic neurotransmission within the mesolimbic and mesocortical systems contributing to anhedonia and loss of motivation in depressive disorders (Chaudhury et al. 2013; Ben Zion et al. 2006; Berridge and Kringelbach 2008). A significant reduction in dopamine transporter (DAT) uptake is observed in anhedonic depressed patients using SPECT imaging (Sarchiapone et al. 2006), as well as increased striatal D2 receptor binding in MDD (D’Haenen and Bossuyt 1994), suggesting decreased dopamine turnover. Similarly, depression in PD has been shown to be associated with both nigral and mesolimbic dopaminergic pathway dysfunction. A reduced volume in the ventral tegmental area (VTA), cingulate and amygdala on MRI, along with reduced [11C]RTI-32 dopamine transporter uptake in limbic regions in PD patients with depression may relate to the loss of dopamine projections from the VTA (Remy et al. 2005) and also loss of substantia nigra (SN) neurones (Fischer et al. 2021). Whilst PD with dementia shows reduced VTA neurone numbers, changes in VTA neurone number in DLB appear to be mild although neurone dysfunction has not been established (Patterson et al 2018). However both DLB and PD show loss of dopaminergic neurones from the SN, and the contribution of SN dopamine depletion is unknown (Patterson et al 2018). What role dopaminergic system plays in DLB patients with depression is unclear.

The aim of this study was to investigate dopaminergic neurotransmission in relation to depression in DLB. We focussed on the subgenual anterior cingulate cortex (sgACC) as a key cortical region important in mood regulation and in the symptomatic expression of depression, and as a region vulnerable to α-synuclein pathology in DLB (Drevets, Savitz, and Trimble 2008; Rodriguez-Cano et al. 2014). Little is known about how changes in dopaminergic neurotransmission in the sgACC might relate to development of depression in DLB, despite the high levels of α-synuclein and other neurodegenerative pathology typically found in the cingulate cortex (Patterson et al. 2018). The sgACC shows changes in cortical and subcortical connectivity based on diffusion tensor imaging in PD (Uhr, Tsolaki, and Pouratian 2022) where depression is also a common feature (Jellinger 2023). We therefore assessed dopaminergic innervation in the sgACC using a combination of stereology-based approaches, high resolution stimulated emission depletion (STED) microscopy, and protein determination in the sgACC using post-mortem tissue samples from DLB cases with and without depression and in normal controls to investigate associations with depression.

## Materials and Methods

### Clinical Cohort

All post-mortem human brain tissue was obtained from the Newcastle Brain Tissue Resource (NBTR). Ethical approval for the study was granted by Newcastle and North Tyneside-1 National Health Service (NHS) Research Ethics Committee. All participants had received clinical assessments during their life and had consented to the use of their brain tissue for research purposes. Neuropathological assessment was according to standardised diagnostic procedures and with clinical data was used to make a clinico-pathological diagnosis (Braak et al. 2006; McKeith et al. 2005; Thal et al. 2002). A total of 17 controls, 15 DLB cases without depression and 13 DLB cases with depression were included in pathological cohort, with 12 from each group used for biochemical analysis. Four DLB cases also fulfilled the neuropathological criteria for high AD neuropathological change and could therefore be classified as neuropathologically mixed DLB/AD with a Lewy body disease (LBD) clinical phenotype (Montine et al. 2012; Walker et al. 2015). All cases were matched as closely as possible for age, sex and post-mortem delay. The inclusion criteria for depression diagnosis was made using the Cornell Scale for Depression in Dementia (CSDD) (score≥8) as a validated rating scale (Alexopoulos et al. 1988). Alternatively, the Geriatric Depression Scale (GDS) (score≥10) was used when the CSDD was not available. This has been shown to have acceptable qualities when applied to demented elderly patients (Korner et al. 2006).

### Immunohistochemistry

Formalin fixed paraffin embedded tissue blocks from the right hemisphere containing the sgACC (Brodmann area: BA25) sampled at the level of the rostrum of the corpus callosum were cut at 10μm using a rotary microtome. Following paraffin wax removal through xylene, sections were rehydrated in decreasing concentrations of ethanol. Citrate buffer heat-induced antigen retrieval (10min, 0.1M, pH 6.0) was used, followed by incubation with 3% hydrogen peroxide for 20 minutes. Tissue sections were incubated with primary antibodies at 4°C overnight (see supplementary Table 1). Following washes in 10mM tris-buffered saline containing 0.1% v/v Tween 20 (TBST) pH 7.6, sections were incubated with horseradish peroxidase (HRP) polymer conjugated Universal Probe for 30 minutes at room temperature, followed by incubation for 30 minutes with HRP reagent (MenaPath, Menarini Diagnostics UK). Visualisation was performed using diaminobenzidine substrate (Menarini X-Cell-Plus HRP Detection Kit; Menarini Diagnostics UK), for two minutes. Sections were counterstained with Mayer’s haematoxylin and mounted using DPX mounting medium (Cell Path, UK). For fluorescent tissue labelling for STED, sections were blocked using 10% normal goat serum (NGS: Sigma G9023) for one hour at room temperature, followed by incubation with the primary antibodies overnight at 4°C. After washes in TBS, sections were incubated with secondary fluorescent antibodies in TBS and 10% NGS for 1 hour at room temperature. To stain nuclei, sections were incubated with TO-PRO™-3 Iodide (Invitrogen, UK) for 15 min and mounted using ProLong Glass Antifade Mountant (Thermo Fisher, UK) using high precision coverslips (0.170 ± 0.01 mm thick; Roth, Germany, LH25.1) to improve image quality.

### Estimation of Pathology Load

Densitometric analysis was used to assess the percentage area stained of immunoreactivity within the region of interest. The images were captured at 10X magnification and imported into Fiji Image J analysis software (Windows 64-bit: https://fiji.sc) for densitometric analysis (Figure 2 D-F). The Red-Green-Blue (RGB) thresholds were adjusted manually for each antibody to eliminate the detection of non-specific background staining. The percentage area stained for each antibody was quantified overall, as well as in cortical layers II, III and V. The mean percentage area stained per case was calculated from the mean values obtained across all images taken.

**Figure 1:**
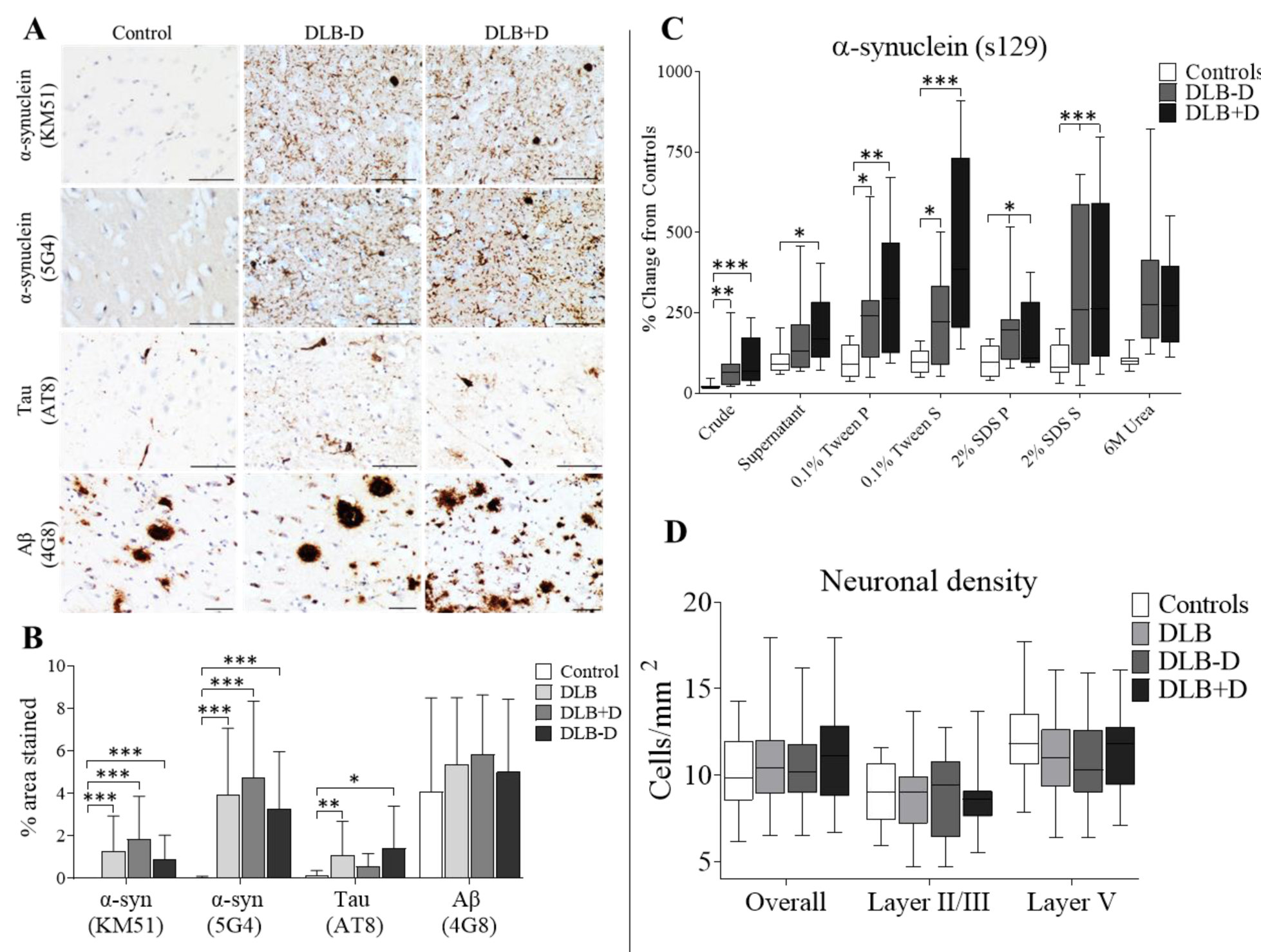
Effect of Pathology on the sgACC. **A)** Photomicrographs of α-synuclein (KM51), α-synuclein (5G4), Tau (AT8) and Aβ (4G8) pathology in sgACC (BA25) in controls, DLB cases with and without depression. Magnification x10; scale bars represent 100μm. **B)** α-synuclein (KM51), (5G4), Tau (AT8) and Aβ (4G8) pathology (% area stained) in sgACC in Controls, DLB cases overall, with and without depression; (\**p*<0.05, \*\**p*<0.01 and *** *p*<0.001, compared to control group). **C)** Dot blotting of α-synuclein phosphorylated at position serine 129 in sgACC within different tissue fractions: crude, supernatant (cytoplasmic soluble proteins), 0.1% Tween pellet (soluble membrane bound proteins), 0.1% Tween supernatant (Tween soluble membrane proteins), 2% SDS pellet (insoluble membrane bound proteins), 2% SDS supernatant (SDS soluble membrane proteins) and 6M Urea supernatant (highly insoluble proteins). **D)** Neurones immunopositive for the neuronal marker HuD were determined in layers II/III and layer V and overall (cells per/mm^2^) within sgACC.

**Figure 2.**
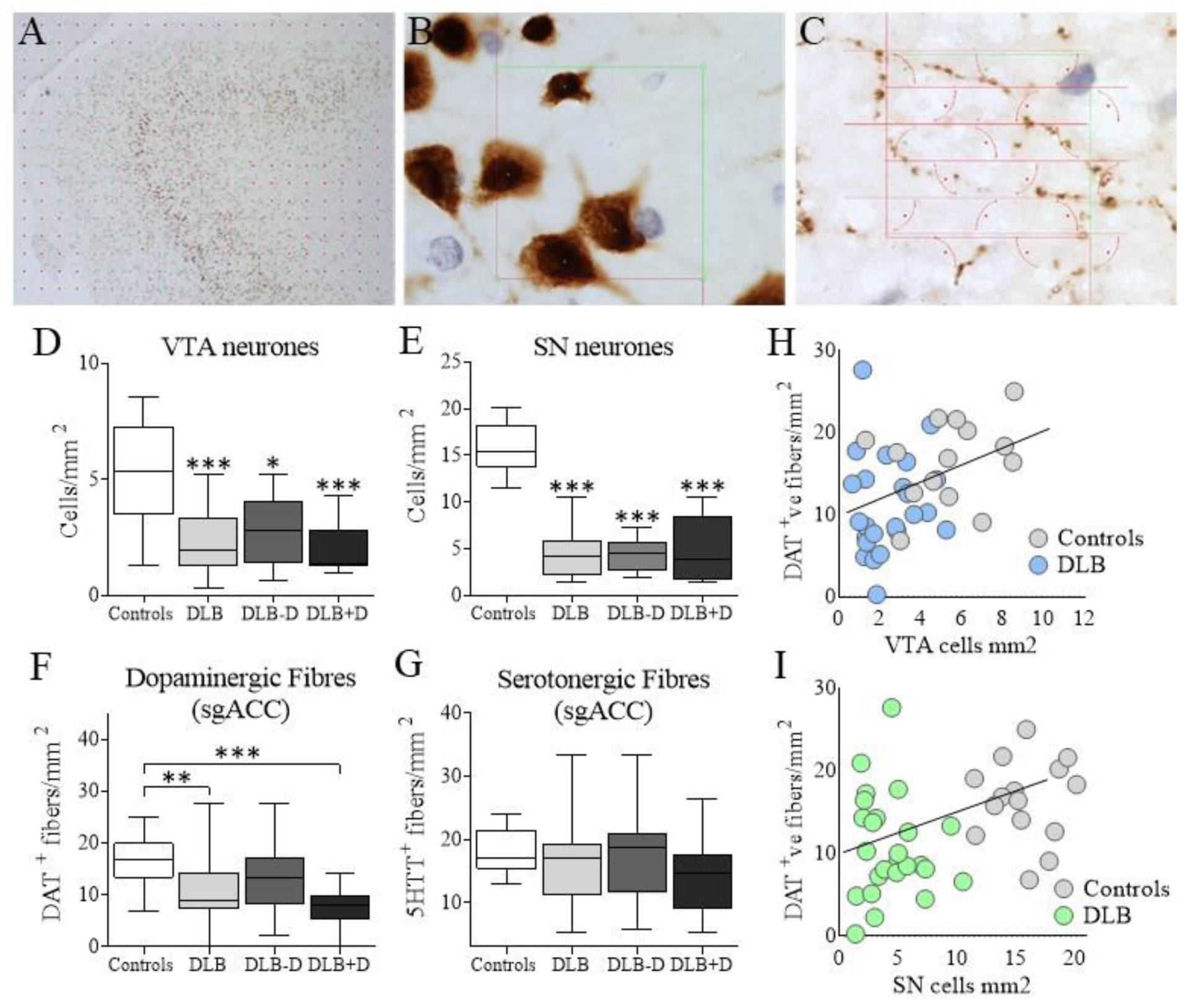
Analysis of Dopaminergic cells in Ventral Tegmental Area and Substantia Nigra, and Dopaminergic and Serotonergic fibres in sgACC. **A)** For stereological imaging, a region of interest was drawn at 1.25X and a point grid was superimposed over the area, with green points representing the coordinates of sampling; **B)** Randomly sampled frames in the x-y axis were taken at 63X, with neurons counted within a dissector frame of known dimensions to allow for an estimation of neurone numbers; **C)** For the estimation of monoaminergic fibres within the outlined reference space, the fibres intersecting (marked green) a grid of randomly orientated cycloids were counted; **D)** pigmented dopaminergic neurones in the VTA; **E)** pigmented dopaminergic neurones in the SN; **F)** Dopaminergic fibres (DAT positive) in sgACC; **G)** Serotonergic fibres (5HTT positive) in sgACC were assessed in controls, DLB cases overall, DLB cases without (DLB-D) and DLB cases with depression (DLB+D) (Significant at \**p*<0.05, \*\**p*<0.01 and \*\*\**p*<0.001); **H)** Correlation analysis between dopaminergic neurones in ventral tegmental area (VTA) (*r*=0.337, *p*=0.007), and **I)** substantia nigra (SN) (*r*=0.313, *p*=0.011) with DAT positive fibres in sgACC.

### Analysis of Neuronal Density

An adapted stereological method was used to estimate the number of neurons within the sgACC. Within each coronal section, a region of interest was drawn at low magnification (1.25X) using the Zeiss Z1 microscope with a motorised stage. A randomly oriented point grid was superimposed over the observed image ensuring the sampling of the structure in a systematic and unbiased manner through its x-y axis. The distance between points on the grid was determined and HuD positive neurons were counted within a disector frame of known dimensions using 63X oil immersion objective. The neuronal density within the sgACC was determined automatically by the Stereologer software (cells per µm^2^), with values converted to cells per mm^2^.

### Estimation of dopaminergic and serotonergic fibre density

Midbrain tissue sections containing the ventral tegmental area (VTA) and substantia nigra (SN) at the insertion of the oculomotor nerve were used to estimate pigmented neurone numbers as previously described (Patterson et al. 2018).

For dopaminergic and serotonergic fibre analysis in sgACC, the images were captured using a Zeiss Z1 microscope with a motorised stage and MRc camera (Zeiss, Germany) coupled to a PC. Stereologer software (Stereologer, Bethesda, MD, USA) was used to ensure adequate and unbiased sampling. The region of interest was drawn at 1.25X magnification and a randomly-oriented point grid was superimposed over the observed image (Figure 2). Within the delineated region of interest 10-15 frames were captured at 10X magnification. Estimation of dopaminergic (dopamine transporter, DAT) and serotonergic (serotonin transporter, 5HTT) positive fibres within sgACC was assessed by counting the number of intersections between the linear probe and lines that represent the surface feature within the dissector frame of known dimensions. The isotropic interaction between the linear probes and the surface feature was achieved through the use of VUR (vertical uniform random) sections in combination with sine-weighted line probes termed cycloids (West 2013).

### STED analysis

Triple-colour stimulated emission depletion (STED) microscopy was applied to assess DAT or 5HTT co-localisation with a presynaptic terminal marker (SNAP25) and phosphorylated α-synuclein (s129) which provides an indication of both physiological and pathological α-synuclein (Ramalingam et al. 2023; Okochi et al. 2000; Fujiwara et al. 2002) in relation to depression in DLB. STED 3D images were acquired using Leica SP8 STED microscope and Application Suite X software (LAS X; Leica Microsystems) with 100x/1.4NA STED white oil immersion objective. Images of 256 × 256 pixels were obtained using 35x optical zoom, resulting in a pixel size of 13 × 13 nm. The gating parameters were set to 1.5-6.0 ns for the 532nm laser, and 0.5-6.0 ns for the 594nm and 647nm lasers. Atto647N was imaged with 645nm excitation, 652-750 emission and 775nm depletion wavelengths, Alexa Fluor 532 with 527nm excitation, 533-581 emission and 660nm depletion wavelengths, whereas Alexa Fluor 594 with 591 excitation, 598-751 emission and 775nm depletion wavelengths. All images were deconvolved using Huygens Essential Software (Scientific Volume Imaging, Netherlands).

Deconvolution settings were optimised to increase the final quality of the image. The number of iterations were set at 40, and the signal to noise ratio at 7. The quality threshold was set at 0.05, which is a restoration parameter that makes deconvolution stop when the change in the quality criterion between two consecutive iterations is below its value. The Object Analyser Advanced tool in Huygens was used to create 3D surfaces for each channel and obtain the quantitative measures of individual particles. A size threshold was applied to remove objects that were too small, with volume set at 10 voxels. The seed and threshold criteria were optimised for individual channels. For the 488nm channel, the seed was set at 5% and threshold at 30%, for 594nm channel, the seed was set at 5% and threshold at 15%, and for 647nm channel, the seed set at 10% and the threshold at 25%. Co-localisation measurements were used to assess spatial overlap between structures in different data channels. Synapses were defined by an overlap of greater than 80% of SNAP25 staining and DAT or 5HTT staining. Alpha synuclein positive synapses were defined where s129 staining was observed to overlap with the synapse exceeded 50% of stained volume.

### Western Blot and Dot Blot analysis

For monoaminergic marker analysis, approximately 50mg unfixed frozen grey matter from the left hemisphere corresponding to sgACC was homogenised using a rotor-stator homogeniser in ice cold lysis buffer consisting of 0.2M triethylammonium bicarbonate (TEAB) pH 7.2 (Sigma-Aldrich, MO, USA), and 1X EDTA free protease inhibitor cocktail (Complete, Roche, UK). Protein concentration in samples was determined using Bradford assay (Bradford, 1976) against standards of known protein concentration prepared using bovine serum albumin (BSA; Sigma-Aldrich, UK).

For phosphorylated α-synuclein analysis, total protein homogenate was fractionated to extract soluble and insoluble proteins (Culvenor et al. 1999). The soluble protein fraction containing supernatant was extracted following centrifugation (20,000 x g for 45min at 4°C), and the pellet re-suspended in 500μl of 0.1% Tween20 in 0.2M TEAB. The soluble membrane bound protein fraction was stored at -30C°. The remaining sample was centrifuged to extract Tween20 soluble membrane bound proteins. The pellet was re-suspended in 500μl of 2% sodium dodecyl sulphate (SDS), and the insoluble membrane bound protein fraction was stored at -30C°. The remaining sample was centrifuged further to extract SDS soluble membrane bound proteins, and the pellet re-suspended in 500μl of 6M Urea. The highly insoluble protein fraction was taken before further centrifugation to extract highly insoluble aggregated proteins. All protein fractions were stored at -30C° prior to use.

Protein samples for dot blot were prepared at 1 µg/µl with 4X Orange G Loading Buffer, 10X NuPAGE Sample Reducing Agent (Invitrogen) and homogenising buffer (0.2M TEAB). Protein standards ranging from 0 μg/μl to 120 μg/μl were prepared using pooled samples from all groups. The samples were denatured at 70°C for 10 minutes. A vacuum-assisted 96-well dot blot apparatus (Hoefer Scientific) was used to blot the samples onto nitrocellulose membrane (Amersham^TM^, 0.2μm NC) using 50µl of sample or standards per well in duplicate. The NC membrane was fixed in 70% methanol for 20 minutes under agitation and blocked using Odyssey blocking buffer (LI-COR), then incubated overnight with primary antibodies diluted in Odyssey blocking buffer and 0.2% Tween20 (Sigma-Aldrich) at 4°C. Following washes, the membranes were incubated with IRDye 800WC secondary antibodies for 1 hour at RT, then incubated with IRDye conjugated GAPDH 680nm (Santa Cruz Biotechnology) for 1 hour at RT. Each membrane was scanned at 700nm (GAPDH) and 800nm (protein of interest) for 2 minutes (LICOR-Fc). The protein intensity bands were quantified using Image Studio Lite (LI-COR). The Lane Normalisation Factor (LNF) was calculated by dividing GAPDH signal for each lane by the highest GAPDH signal. The signal for the protein of interest was then divided by the LNF to normalise the protein of interest.

For SDS polyacrylamide gel electrophoresis (SDS-PAGE), denatured (70°C for 10min) protein samples (1 µg/µl) were loaded onto a NuPAGE 4-12% Bis-Tris Gel (Invitrogen) along with Chameleon 800 pre-stained protein ladder (LI-COR). The gel was electrophoresed in 1X NuPAGE MOPS SDS running buffer with antioxidant (Invitrogen) at 120V for 20 minutes, followed by 160V for one hour. The transfer of proteins to a nitrocellulose membrane was performed using an iBlot2 device (Invitrogen) at 20V for 1min, 24V for 4min and 27V for 5min. Blotted proteins were determined as previously.

### Statistical analysis

Statistical analyses were performed using SPSS Statistics version 22.0. The normal distribution across samples was assessed using Shapiro–Wilk test, with homogeneity of variance determined using Levene’s test. If the assumptions of normality were met, analysis of variance (ANOVA) was used to analyse the data sets between the groups, followed by Bonferroni post-hoc analysis to correct for multiple comparisons. A paired-samples *t*-test was used for pairwise comparisons within groups. Where normal distribution criteria was not fulfilled, a non-parametric Kruskal-Wallis test was used to compare multiple groups, with adjusted *p*-values to correct for multiple comparisons, so that the overall error rate remained at 5%. Friedman’s ANOVA was used for pairwise comparisons within groups. Correlation analyses were carried out using Spearman’s correlation coefficient ρ (rho). Linear discriminant analysis was used to investigate whether separation of groups could be defined on the basis of any of the monoaminergic protein markers.

## Results

### Pathology in Subgenual Cingulate

A significant main effect of diagnosis on α-synuclein (KM51) pathological burden was observed in sgACC *H*(3)=31.370, *p*<0.001, with significant increase identified in DLB cases overall, with or without depression compared to controls (*p*<0.001; Figure 1B). Alpha-synuclein (5G4) pathological burden was also significantly different in sgACC between the groups *H*(3)=40.174, *p*<0.001, with significant increase in DLB cases overall, with or without depression compared to controls (*p*<0.001; Figure 1B). Tau pathological burden was also significantly different in sgACC between the groups *H*(3)=12.140, *p*=0.007. DLB cases with depression did not show significantly higher p-Tau burden in sgACC compared to controls, however DLB cases overall (*p*=0.010) and without depression (*p*=0.015) showed significant elevation of tau burden in sgACC compared to controls. Aβ burden was not significantly different between the groups in sgACC *H*(3)=3.266, *p*=0.352 (Figure 1B).

### Biochemical Analysis of α-synuclein

An antibody to phosphorylated α-synuclein was used to assess biochemical changes in the sgACC using fractionated tissue (Delic et al. 2018). The biochemical analysis of α-synuclein pathological burden in sgACC generally supported immunohistochemical data, and showed no significant difference in α-synuclein levels in sgACC between DLB cases with and without depression. Increased phosphorylated α-synuclein was found in the crude sample *(H*(2)=16.826, *p*<0.001), supernatant (*H*(2)=7.115, *p*=0.029), 0.1% Tween pellet (*H*(2)=11.988, *p*=0.002), 0.1% Tween supernatant (*H*(2)=16.122, *p*<0.001), 2% SDS pellet (*H*(2)=6.504, *p*=0.039), 2% SDS supernatant (*H*(2)=9.159, *p*=0.028) and urea fraction of tissue homogenates (*H*(2)=20.523, *p*<0.001) were seen. DLB cases with depression showed significantly higher s129 α-synuclein burden compared to controls in crude (*p*=0.001), supernatant (*p*=0.029), 0.1% Tween pellet (*p*=0.003), 0.1% Tween supernatant (*p*<0.001), 2 % SDS supernatant (*p*=0.020) and urea (*p*=0.001). DLB cases without depression showed higher s129 α-synuclein immunoreactivity compared to controls in the crude (*p*=0.002), 0.1% Tween pellet (*p*=0.045), 2% SDS pellet (*p*=0.010) and 2% SDS supernatant (*p*=0.035) and 6M Urea tissue fraction (p<0.001). There was no significant difference between DLB cases with or without depression in s129 immunoreactivity (Figure 1C).

The neuronal specific marker (HuD) raised against amino acids 1-300 of human HuD (Szabo et al. 1991) was used to assess neuronal cell density within sgACC between the groups. There was no significant difference observed in neuronal density overall *F*(2, 48)=0.393, *p*=0.677, or within layers II/III *F*(2, 48)=0.018, *p*=0.982, or layer V *F*(2, 48)=1.183, *p*=0.315 of the sgACC between groups (Figure 1D).

### Dopaminergic and Serotonergic Fibres in Subgenual Cingulate

The sgACC receives dopaminergic afferent projections from the VTA with additional input from the SN (Freedman, Insel, and Smith 2000; Vergani et al. 2016; Porrino and Goldman-Rakic 1982) and therefore we assessed the number of pigmented dopaminergic neurons in the VTA and SN, as well as DAT and 5HTT positive fibres in the sgACC between controls and DLB cases overall, and in DLB cases with and without depression. The number of neurons in the VTA was significantly different between the groups *H*(3)=19.056, *p*<0.001, with significantly lower neuronal count observed in DLB group overall (*p*=0.001), DLB cases with depression (*p*=0.001) and DLB cases without depression compared to controls (*p*=0.035; Figure 2D). The number of neurons in the SN was also significantly different between the groups *H*(3)=34.179, *p*<0.001, with significantly lower neuronal count observed in all the groups compared to controls (*p*<0.001; Figure 2E). There were no significant differences in neurone counts between depressed and non-depressed donors in either the VTA or SN.

The number of DAT positive dopaminergic fibres in the sgACC was significantly different between the groups *F*(3, 69)=7.029, *p*<0.001, with significantly lower fibre density observed in DLB cases overall (*p*=0.004) and in DLB cases with depression compared to controls (*p*<0.001) but not in non-depressed DLB donors (Figure 2F). Since serotonergic neurotransmission abnormalities are implicated in the neurobiology of depression, serotonergic innervation of sgACC was assessed using an identical approach. Serotonergic fibres were more abundant in the sgACC compared to dopaminergic fibres χ2(1) = 8.138, *p* = 0.004, however, no significant difference in the number of serotonergic fibres in the sgACC was observed between groups *F*(2, 39)=1.694, *p*=0.197 (Figure 2G). No significant correlations were observed between dopaminergic neurons in VTA or SN, and DAT positive fibre density in sgACC within the groups (supplementary Figure 1). In the combined control and DLB groups however, a significant correlation was observed between DAT positive fibre density and VTA neurones *r*=0.337, *p*=0.007, as well as dopaminergic neurons in SN *r*=0.313, *p*=0.011 (Figure 2H-I).

### Dopaminergic and serotonergic synapses in sgACC

The morphometric findings were extended by assessment of serotonergic (5HTT and SNAP25 positive) and dopaminergic synapses (DAT and SNAP25 positive) in relation to phosphorylated α-synuclein (s129) in the sgACC using STED microscopy (Figure 3). Due to the high resolution analysis provided by STED microscopy, a strictly unbiased stereological approach to image analysis was not possible.

**Figure 3.**
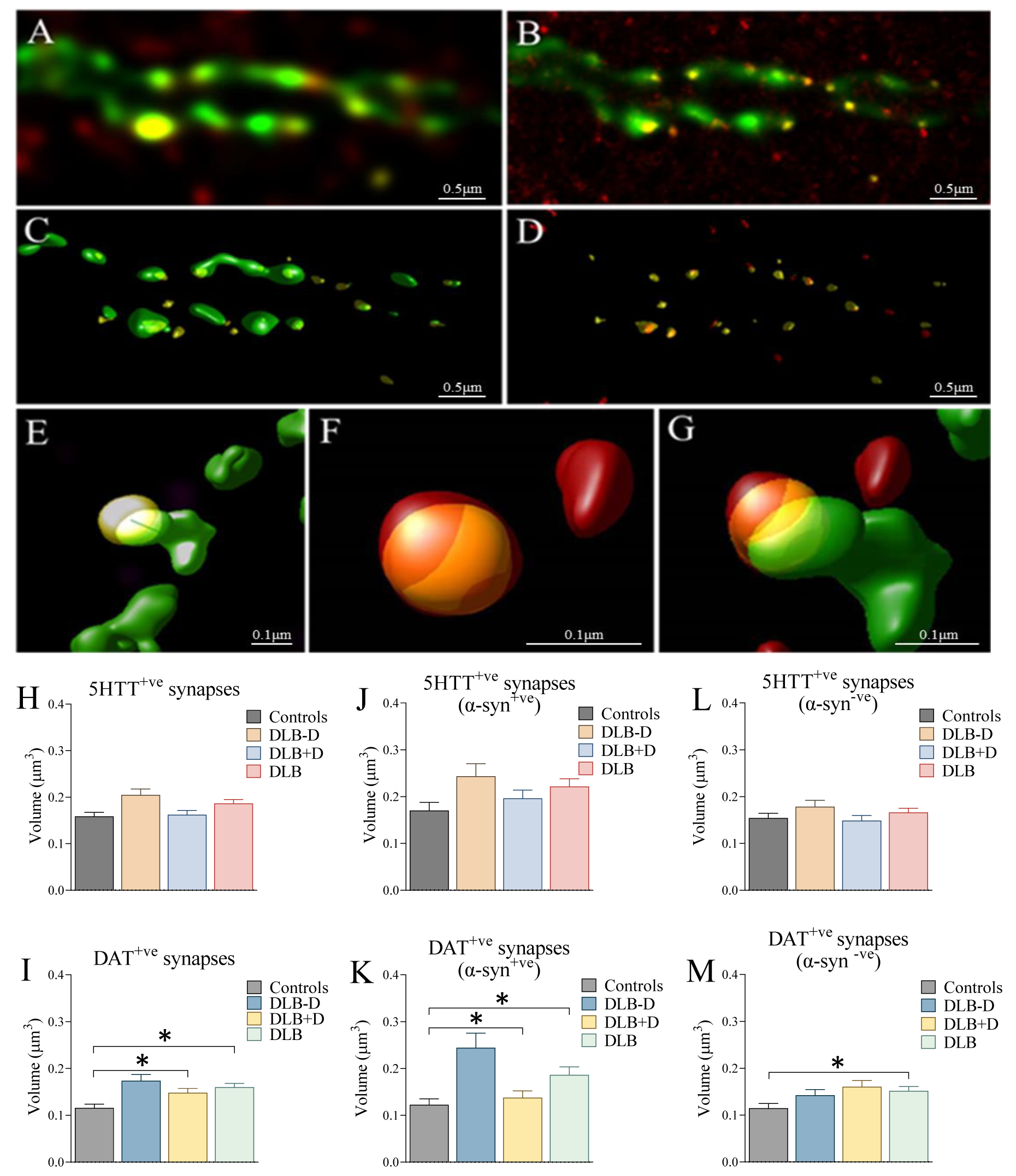
Stimulated Emission Depletion (STED) imaging of Dopaminergic and Serotonergic Synapses in Subgenual anterior cingulate. **A)** - STED 3D image of dopaminergic fibres and terminals (DAT; green), presynaptic terminals (SNAP25; yellow) and phosphorylated α-synuclein (s129; red); **B) -** Deconvolved STED image; **C, E) -** 3D surfaces of dopaminergic fibres and presynaptic terminals; **D**, **F) -** 3D surfaces of presynaptic terminals and α-synuclein; **G) -** 3D surfaces of co-localisation of all 3 channels; **H)** Co-localisation of serotonergic (5HTT +ve) and **I)** dopaminergic (DAT +ve) synapses with pre-synaptic terminal marker (SNAP25) (>80% co-localisation); **J, K)** Serotonergic and dopaminergic synapses α-synuclein (s129) +ve, and **L, M)** α-synuclein (s129)–ve synapses in sgACC were assessed in controls, DLB cases overall, DLB cases without (DLB-D) and DLB cases with depression (DLB+D) (Significant at \**p*<0.05).

No significant difference in the proportion of presynaptic serotonergic terminals *F*(2, 22)=1.258, *p*=0.299 or the proportion of serotonergic synapses containing α-synuclein was observed between the groups *F*(2, 22)=0.057, *p*=0.944. 5HTT containing synapses were not significantly different in size between groups *H*(3)=0.833, *p*=0.842. No significant difference was observed in α-synuclein positive *H*(3)=2.089, *p*=0.554, or α-synuclein negative *H*(3)=0.803, *p*=0.749 serotonergic synapses between the groups (Figure 3H, J, L).

No significant differences were observed in the proportion of dopaminergic synapses (*p*=0.582), or dopaminergic synapses containing s129 between the groups (*p*=0.823). The volume of DAT positive terminals was however, significantly different between the groups *H*(3)=12.007, *p*=0.007, with larger dopaminergic synapses observed in DLB cases overall (*p*=0.011), as well as DLB cases without depression (*p*=0.017) compared to control (Figure 3I). Specifically, DAT synapses containing s129 alpha-synuclein were larger compared to controls in the DLB group overall (*p*=0.043), and in the non-depressed DLB group (*p*=0.016), but not in the depressed DLB group. Alpha-synuclein negative dopaminergic synapses were significantly larger in DLB cases with depression (*p*=0.028) compared to control α-synuclein negative synapses (Figure 3K, M).

Alpha-synuclein is known to interact with synaptic vesicle proteins (Burre et al. 2010; Burre, Sharma, and Sudhof 2014), therefore we determined if the presence of α-synuclein within dopaminergic or serotonergic synapses was related to synaptic activity using SNAP25 as a marker. A significant positive correlation between the volume (μm^3^) of s129 and SNAP25 within presynaptic 5HTT positive terminals was observed in DLB cases overall (*r*=0.288, *p*=0.012), DLB cases without (*r*=0.313, *p*=0.016) and DLB cases with depression (*r*=0.316, *p*=0.030; Figure 4). A significant positive correlation was also observed between the volume of s129 α-synuclein and SNAP25 within presynaptic DAT positive terminals in DLB cases with depression (*r*=0.379, *p*=0.007; Figure 4), but not in DLB cases without depression or controls.

**Figure 4.**
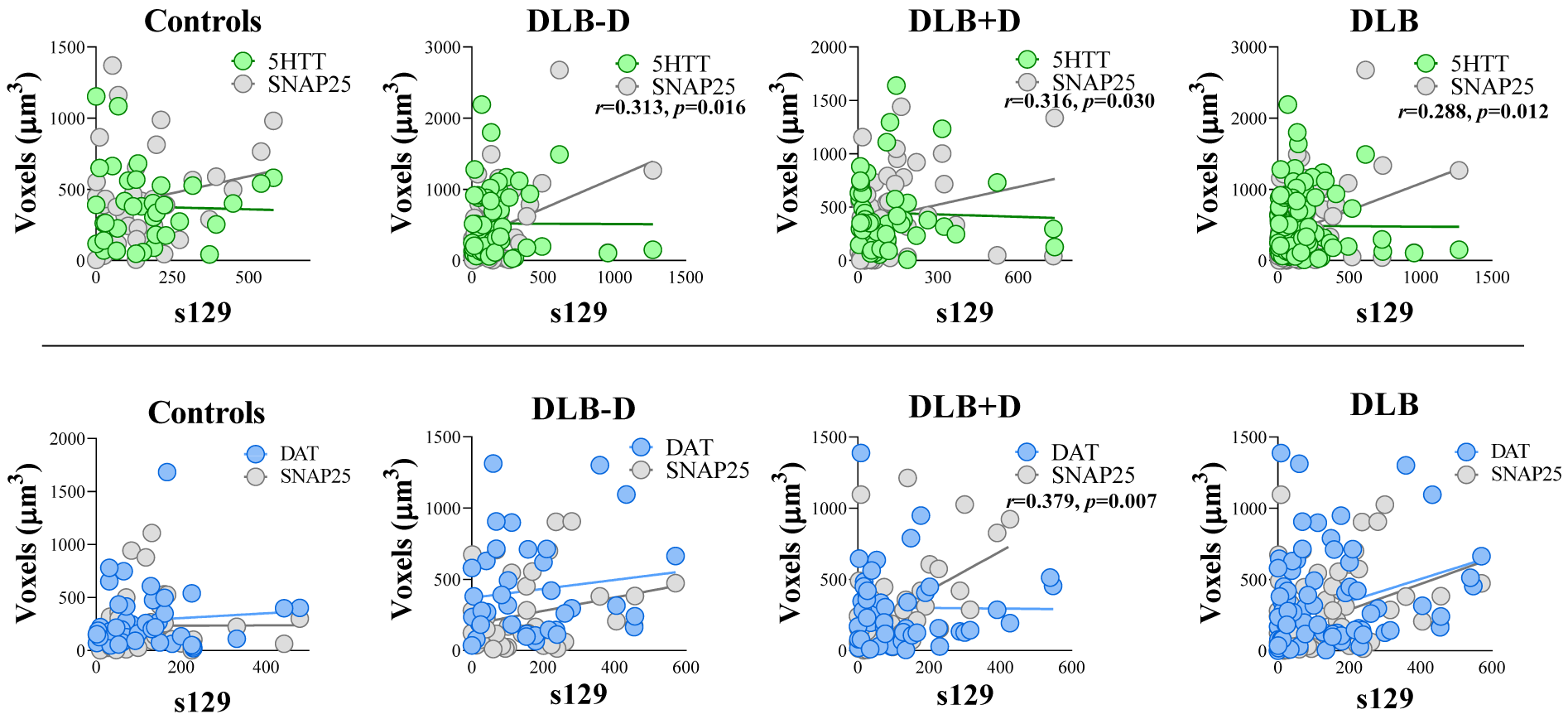
Effect of phosphorylated α-synuclein (s129) on serotonergic and dopaminergic synapses. **Top)** The relationship between 5HTT and SNAP25, and **Bottom)** DAT and SNAP25 with s129 within synapses in the sgACC was assessed using STED microscopy and Spearman’s correlation analysis in controls, DLB cases overall, DLB cases with (DLB+D) and without (DLB-D) depression.

A significant correlation was observed between the number of dopaminergic neurons in VTA and the volume of DAT positive synapses in sgACC in DLB cases with depression (*r*=0.710, *p*=0.049; supplementary Figure 2). Furthermore, a significant correlation was observed between the number of dopaminergic neurons in VTA and the volume of DAT positive synapses in sgACC which were α-synuclein positive in DLB cases with depression (*r*=0.732, *p*=0.016; supplementary Figure 3). No significant correlations was observed between the number of dopaminergic neurons in SN and the volume of DAT positive synapses in sgACC within the groups (supplementary Figure 2).

### Monoaminergic Protein Analysis

To determine the impact of DLB on monoamine neurotransmission we used western and dot blot with specific antibodies to dopaminergic and serotonergic markers. Dopamine transporter (DAT) *F*(3, 56)=4.346, *p*=0.008, tyrosine hydroxylase (TH) *F*(3, 56)=8.762, *p*<0.001, dopamine decarboxylase (DDC) *F*(3, 56)=3.660, *p*=0.018 and dopamine D3 receptor (D3) levels *F*(3, 56)=2.913, *p*=0.042 were significantly different between the groups in the sgACC (Figure 5 A-F). DLB cases overall showed significantly lower levels of DAT (*p*=0.022), TH (*p*=0.002) and DDC (*p*=0.024) compared to controls. Significantly lower levels of DAT (*p*=0.009), TH (*p*=0.003), DDC (*p*=0.041) and D3 (*p*=0.049) were observed in DLB cases with depression compared to controls. DLB cases without depression showed significantly lower levels of TH (*p*=0.012) compared to controls. No significant difference was observed in the levels of dopamine D2 receptors *F*(3, 56)=0.620, *p*=0.605 or D4 receptors *F*(3, 56)=2.115, *p*=0.109 in the sgACC between the groups.

**Figure 5.**
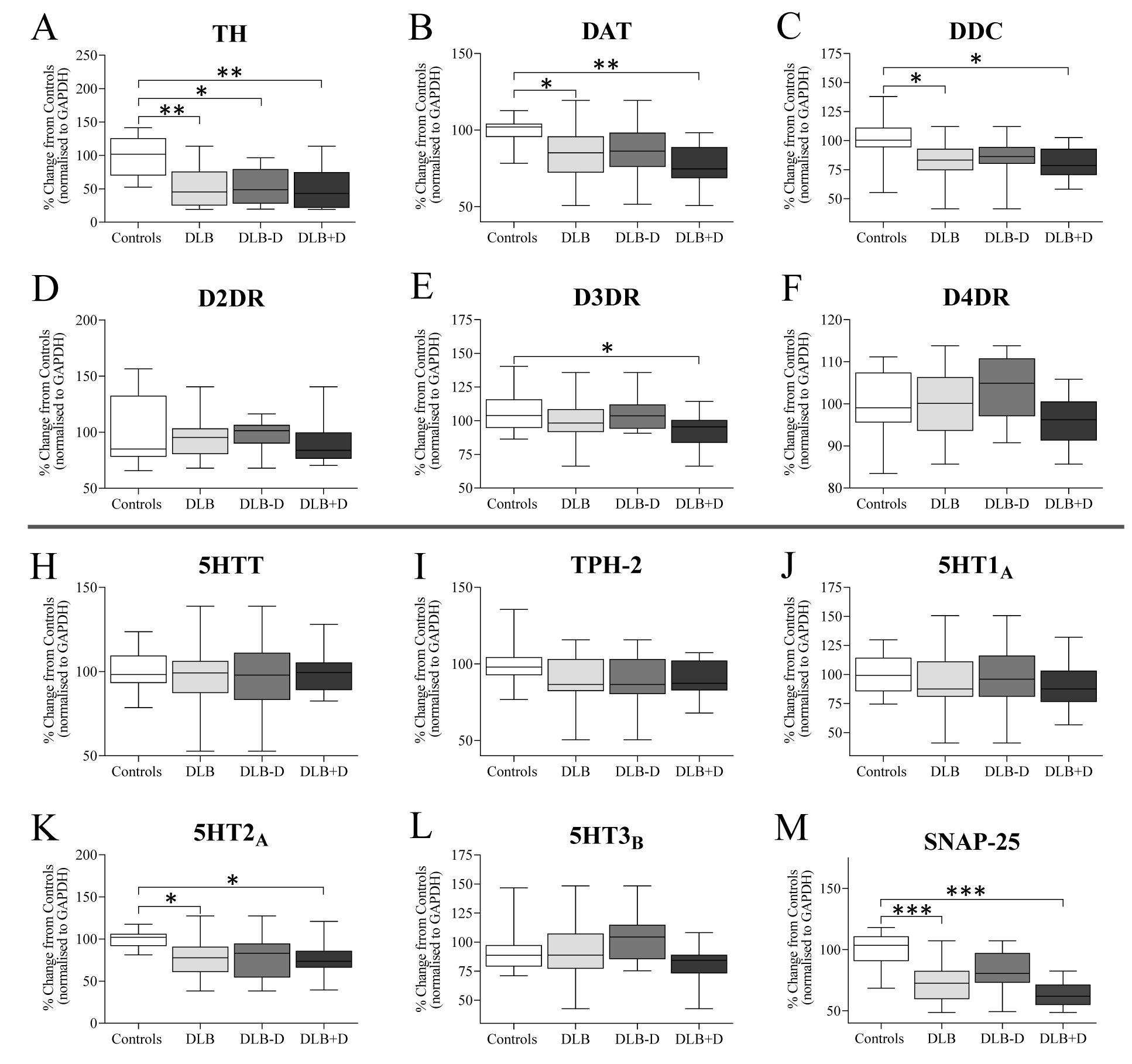
Dot blot analysis of dopaminergic and serotonergic markers and receptors in sgACC. Dopaminergic markers, including **A**) tyrosine hydroxylase (TH), **B**) dopamine transporter (DAT), **C)** dopamine decarboxylase (DDC), and **D-E)** dopamine receptors D2, D3 and D4, as well as serotonergic markers, including **H)** serotonin transporter (5HTT), **B)** tryptophan hydroxylase-2 (TPH-2), and **J-L)** serotonin receptors 5HT1A, 5HT2A and 5HT3B were assessed in sgACC in age matched controls, DLB cases overall, DLB cases with (DLB+D) and without (DLB-D) depression; (\**p*<0.05, \*\**p*<0.05 and \*\*\**p*<0.001, compared to appropriate group).

No significant changes in 5HTT *F*(3, 56)=0.064, *p*=0.978, tryptophan hydroxylase 2 (TPH-2) *F*(3, 56)=1.719, *p*=0.173, 5HT1_A_ receptor *F*(3, 56)=0.258, *p*=0.855 and 5HT3_B_ receptor *F*(3, 56)=2.156, *p*=0.062 was observed between the groups (Figure 5 H-L). A significant difference in 5HT2_A_ receptor protein levels was observed in the sgACC between groups *F*(2, 34)=3.858, *p*=0.019, with significantly lower 5HT2_A_ receptor protein levels observed in DLB cases overall (*p*=0.026) and in DLB cases with depression compared to controls (*p*=0.044).

Heat map analysis showed no clear separation of disease groups based on monoaminergic proteins levels (Figure 6). A linear discriminant analysis was used to separate the cases based on monoaminergic protein levels to determine the greatest influence on depression status. DAT (Wilks’ Lambda, 0.535, p=0.0004) and D4DR (Wilks’ Lambda 0.442, p=0.0002) in combination showed best predictive values in separating the groups, showing 78% accuracy in re-classifying the original data.

**Figure 6.**
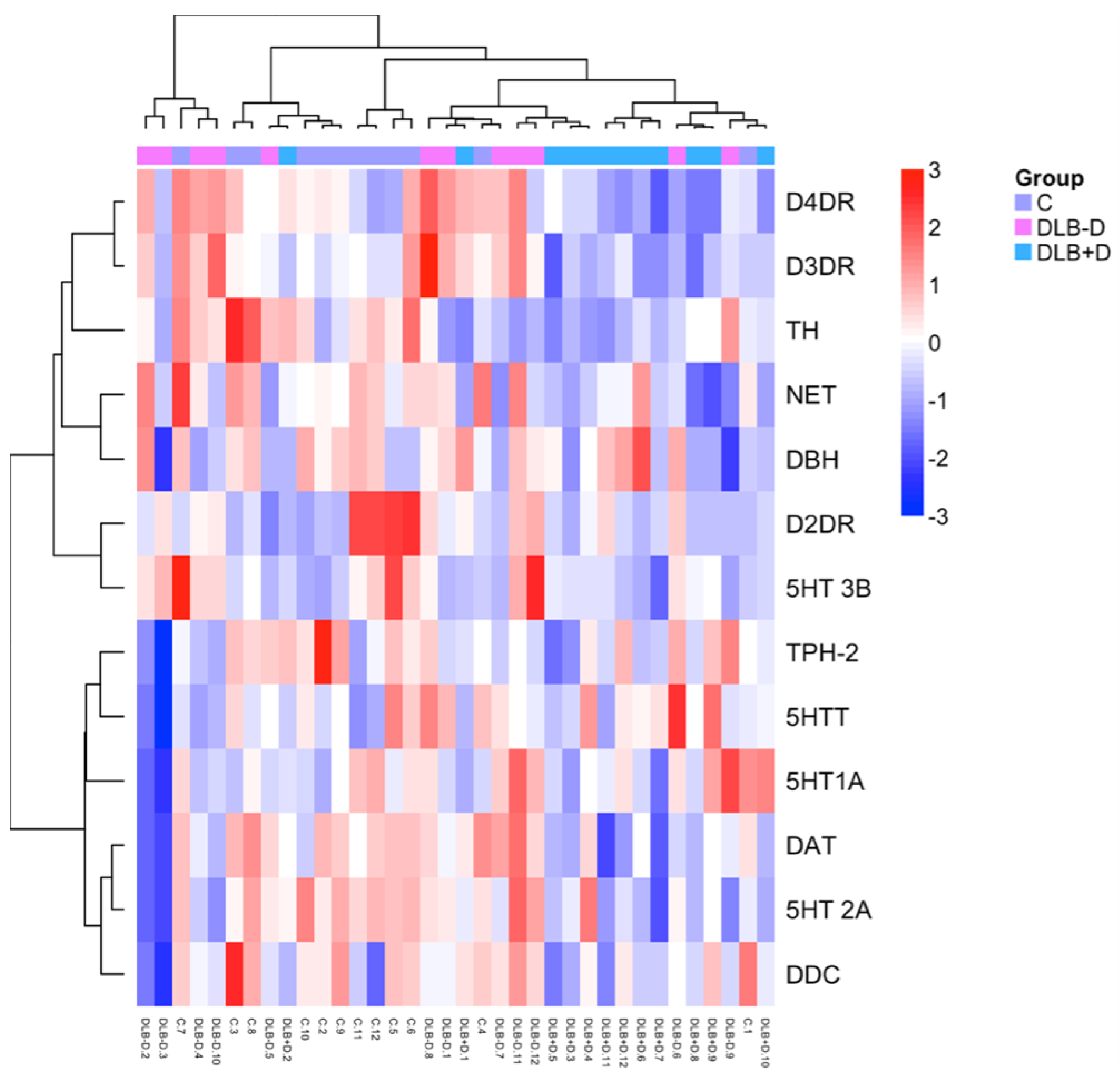
Heat map analysis of monoaminergic markers and receptors in the sgACC. Hierarchical clustering analysis was performed based on monoaminergic protein relative expression in sgACC in controls, DLB cases with (DLB+D) and without (DLB-D) depression.

## Discussion

Individuals with DLB are at high risk of developing major depression as part of the syndrome, with depression being present from an early stage (Fujishiro et al. 2015) with a significant impact on quality of life (Gatt et al. 2017; McKeith et al. 2017). Depression in DLB correlates with α-synuclein pathology and Lewy bodies in late onset MDD found in older populations, indicating that α-synuclein pathology may be a significant factor in development of MDD in older individuals (Nunes et al. 2022). The sgACC is an area intimately involved in the aetiology of depression and shows high α-synuclein pathological burden in DLB (Patterson et al. 2018), with known structural and functional changes observed in early onset major depression (Drevets, Savitz, and Trimble 2008; Rodriguez-Cano et al. 2014). In this study despite high levels of α-synuclein in the sgACC, there was no significant difference between depressed and non-depressed DLB donors for α-synuclein pathology burden suggesting pathology within the sgACC is not a significant driver of depressive symptoms. Major neurone loss within the sgACC in DLB does not seem to occur, and there is no significant association with depression (see Figure 1). However, little is known how changes within monoaminergic, and particularly serotonergic and dopaminergic, neurotransmission in the sgACC modulates development of depression in DLB. Although the underlying basis for depression in DLB is complex, our results based on monoaminergic fibre analysis suggest that there is reduced dopaminergic innervation in DLB without a similar change in serotonergic transmission. This reduced dopaminergic innervation in sgACC, with decreased levels of dopaminergic markers and changes in dopaminergic synaptic function, may be a contributor to depression in DLB. Clinical trials of D2 dopamine receptor agonists, such as pramipexole and pergolide have shown the efficacy in reducing depression in patients with Parkinson’s disease (Rektorová et al. 2003; Ziaei, Ardestani, and Chitsaz 2022; Seppi et al. 2011), as well as symptoms of major depression in patients without Parkinson’s disease (Whitton et al. 2020; Lattanzi et al. 2002).

The mesolimbic dopamine pathway sends projections from the VTA to widespread cortical and subcortical regions including the cingulate cortex (Porrino and Goldman-Rakic 1982), although projections from the dorsal tier of the SN also project to ventral striatal (Lynd-Balta and Haber 1994a, 1994b), thalamic (Melchitzky, Erickson, and Lewis 2006), and cingulate (Porrino and Goldman-Rakic 1982). VTA dopaminergic neurons play a role in reward and stress (Pignatelli and Bonci 2015) with dysfunction of dopaminergic neurotransmission in the mesolimbic system contributing to anhedonia and loss of motivation in depressive disorders (Chaudhury et al. 2013). In this study we found a decrease in VTA cell number in DLB cases overall, but no significant loss in relation to depression in DLB. Compared to the major loss of neurones in the SN (Patterson et al. 2018), the VTA shows relative preservation of neurones in PD (Alberico, Cassell, and Narayanan 2015; McRitchie, Cartwright, and Halliday 1997; Patterson et al. 2018). This may be due to expression of Calbindin by VTA dopaminergic neurons, since Calbindin expression is suggested to be neuroprotective for dorsal tier SN dopaminergic neurones (Pan and Ryan 2012; Pereira Luppi et al. 2021). Studies have shown that in PD more severe cell loss and gliosis is associated with the presence of depression (Paulus and Jellinger 1991), with neuronal density in the VTA and to a lesser extent SN having a significant effect (Wilson et al. 2013), in line with the current findings. In Lewy body disease (PD, PDD, and DLB), higher midbrain α-synuclein burden associates with depressive symptoms (Patterson et al. 2018). This suggests that mesolimbic and mesocortical projections from the midbrain rather than nigrostriatal motor dopaminergic changes contribute to development of depression in DLB, as seen in PD (Wang, Zhao, and Schrag 2023).

In this study, we observed moderate dopaminergic innervation in all cortical layers in the sgACC and reduced dopaminergic fibre density in sgACC in DLB cases with depression compared to controls, which may be an indicator of the reduced innervation from the VTA. In PD, motor symptoms appear when around 50% of SN dopaminergic neurons are lost (Ross et al. 2004), whereas the extent of striatal innervation at the onset of motor symptoms is more severe, showing about 80% dopamine loss (Bernheimer et al. 1973). Dopaminergic axons and terminals are the main site of pathology, with the neurodegenerative process in PD thought to follow a retrograde pathway originating in the striatal terminals (Hornykiewicz 1998). The enhanced dopaminergic fibre loss seen in this study in DLB cases with depression in the absence of severe (∼20%) VTA neuronal loss may represent a similar effect with axon and terminal loss prior to major cell death.

Our STED based analysis showed phosphorylated α-synuclein within dopaminergic synapses in the sgACC and increased phosphorylated α-synuclein in sgACC tissue homogenates. In depressed DLB cases, the presence of synaptic α-synuclein showed a positive correlation with SNAP25, where elevated α-synuclein corresponded with elevated SNAP25, a finding also seen with serotonergic terminals. One possibility is that pathological α-synuclein causes sequestration of SNAP25 within synapses leading to synaptic dysfunction. Alpha-synuclein is a presynaptic protein that modulates synaptic activity through effects on vesicle recycling and release (Burre, Sharma, and Sudhof 2014; Burre et al. 2010; Diao et al. 2013; Ramalingam et al. 2023). Endogenous α-synuclein in particular, promotes synaptic vesicle generation by assisting with the formation of SNARE complexes (Burre, Sharma, and Sudhof 2014; Chandra et al. 2005; Sun et al. 2019). The positive correlation identified in the current study may represent a pathophysiological response of synapses to increase activity and maintain normal neurotransmission following reduction in cell numbers (Reeve et al. 2018). Pathological fibrillar α-synuclein can however, rapidly promote aberrant synaptic activity following application to neurones (Volpicelli-Daley et al. 2011). In forming aggregates within synapses, fibrillar α-synuclein may lead to an effective depletion of the endogenous pool of functional SNARE proteins including SNAP25, reducing formation of a functional synaptic vesicle pool.

Fibrillar and oligomeric α-synuclein also has direct effects on synaptic machinery with further depletion of SNARE complexes (Choi et al. 2013; Larson et al. 2017; Rockenstein et al. 2014). This may underscore the observed increase in SNAP25 seen within α-synuclein containing dopaminergic synapses, with SNAP25 increased, but non-functional due to sequestration in α-synuclein aggregates (Choi et al. 2018). The increased volumes of α-synuclein negative synapses in depressed DLB donors may be a compensatory response to reduced functional activity of α-synuclein containing synapses in depressed cases. The combined effects of dopaminergic synapse reduction due to cell loss and reduced synaptic efficiency in remaining synapses may cause an effective depletion of dopamine to the sgACC cortex and contribute to depressive symptoms.

In this study, we found a decrease in dopaminergic proteins in DLB cases with depression compared to controls, including DAT, TH and DDC. Using hierarchical clustering, DAT and DRD4 showed the largest associations with depression in DLB (see Figure 6). These reductions align with DAT fibre loss in depressed DLB cases and reinforce the idea that dopaminergic innervation is lost in DLB generally, but particularly in DLB experiencing depression (Paulus and Jellinger 1991; Fischer et al. 2021; Patterson et al. 2018). Dopaminergic deficits are implicated in depression (Lambert et al. 2000) with specific antidepressants able to enhance mesolimbic dopaminergic neurotransmission (Maj, Wedzony, and Klimek 1987). Reduced striatal DAT binding has been observed in depressed patients with anhedonia (Sarchiapone et al. 2006), as well as in PD and DLB, with DLB cases showing more severe DAT loss in caudate compared to PD (Walker et al. 2004) along with reduced ACC DAT binding in DLB (Marquie et al. 2014). The use of dopamine agonists in treatment resistant depression is shown to have significant benefits and may have possible benefits in DLB if a suitable treatment regime can be found (Tundo et al. 2023; Tundo, de Filippis, and De Crescenzo 2019).

Our results indicate minimal changes in post-synaptic D2 sgACC receptors in DLB cases with depression indicating that compensatory effects as seen in the striatum do not occur in the sgACC and that other dopaminergic receptor changes may occur. The highest levels of D2 dopamine receptors are expressed in the striatum, nucleus accumbens and olfactory tubercle, with significant levels in the SN, VTA, hypothalamus, amygdala, hippocampus and cortical areas (Gurevich and Joyce 1999). In PD, progressive loss of SN dopaminergic neurons results in reduced striatal dopamine and an increase in striatal D2 receptor density as a compensatory mechanism in response to low dopamine (Rinne et al. 1995). Increased dopamine D2/D3 receptor binding and lower DAT activity has also been shown in MDD, potentially reflecting compensatory changes due to the impairment in mesolimbic dopaminergic neurotransmission (Klimek et al. 2002; Allard and Norlen 2001). This does not appear to occur in DLB however, where reduced striatal and cortical D2 receptors occurs (Piggott et al. 1999; Piggott et al. 2007).

D3 receptors (DRD3) are expressed on postsynaptic and presynaptic neurons and have the highest affinity for dopamine (Sokoloff et al. 2006). DRD3 receptor expression is highest in limbic areas such as the amygdala and nucleus accumbens (Sokoloff et al. 1992) with lower levels in cortical regions where postsynaptic DRD3 expression is principally on a subset of glutamatergic neurons (Li and Kuzhikandathil 2012; Lidow et al. 1998; Clarkson et al. 2017). Activation of cortical DRD3 receptors on glutamatergic neurones results in enhanced glutamate release (Lopez-Gil et al. 2007; Lorrain et al. 2003; Moghaddam et al. 1997). The DRD3 is a clinically relevant target for depression treatment following identification of DRD3 downregulation in MDD (Leggio et al. 2013), and dopamine D3 (and D2) receptor agonists such as pramipexole and ropinirole have shown antidepressant and anxiolytic effects in humans and animals (Tadori et al. 2011; Zarate et al. 2004; Rogoz, Skuza, and Kllodzinska 2004; Leggio et al. 2013; Leentjens et al. 2009; Seppi et al. 2011). A reduction in DRD3, particularly in the ventral striatum has been observed in PD cases (Joyce 1993), but with variable findings of either downregulation (Sweet et al. 2001) or no change (Piggott et al. 1999) in DLB. Other studies have shown elevated DRD3 binding in non-demented PD cases, with reduced DRD3 binding in PD cases with dementia (Joyce et al. 2001), suggesting that DRD3 may modulate cognitive function (Sharp et al. 2020). The current results suggest that reduced DRD3 levels may play a role in depression in DLB and DRD3 could be targeted through use of specific dopaminergic agents. Certainly, pramipexole and potentially ropinirole improve depressive symptoms in small scale trials in PD and may provide benefit for depression in DLB, although careful dosage and monitoring may be needed to prevent unwanted side effects including impulse control disorders but importantly hallucinations (Tundo et al. 2023; Tundo, de Filippis, and De Crescenzo 2019; Gencler, Oztekin, and Oztekin 2022; Leentjens et al. 2009; Roy et al. 2018; Seppi et al. 2011). Effects on cognition may also occur with studies in PD indicating improved working and episodic memory with dopamine agonists in certain cases, indicating that dopaminergic agonists may be useful (Costa et al. 2009; Roy et al. 2018).

Our findings show no significant changes in sgACC DRD4 levels in DLB although an association based on hierarchical clustering. The DRD4 has the lowest level of expression of dopamine receptors in the brain, with greatest densities in anterior regions of the limbic and cortical forebrain, suggesting a function in cognitive, executive and motivational regulation (Rondou, Haegeman, and Van Craenenbroeck 2010). In line with previous studies showing that DRD4 are not implicated in the modulation of depressive-like behaviours (Basso et al. 2005), our findings suggest that DRD4 may not be a treatment target.

In contrast to dopaminergic changes, we found limited evidence for serotonergic involvement in depression in DLB, comparable with similar studies (Mizutani et al. 2022; Wilson et al. 2013). Serotonergic neurotransmission has been suggested to be severely impaired in DLB compared to AD patients with depression, with reduced 5HT and 5HT-metabolite concentrations in the prefrontal cortex, temporal lobe, limbic regions, occipital cortex and hippocampus (Vermeiren et al. 2015). The dorsal raphe shows LB pathology in DLB and PD, with a suggested loss of serotonergic innervation to the forebrain (Ballard et al. 2013). Studies have however shown variable cell loss with some studies showing reduction (Seidel et al. 2015), or no change (Cheshire et al. 2015; Mizutani et al. 2022; Halliday et al. 1990) within the dorsal raphe in PD or PDD. In one report in DLB, neuronal loss is present within both the dorsal and median raphe (Benarroch et al. 2007). In the current study, due to the complexity of the dorsal raphe anatomy, tissue sampling strategy, and differences in dorsal raphe projection sites (Ren et al., 2018), dorsal raphe neurone numbers were not assessed. We however found no change in sgACC serotonergic fibre density in DLB indicating neuronal loss in DLB, similar to the findings of unaltered fibre density in the amygdala and dorsal prefrontal cortex in LBD donors with and without depression (Mizutani et al. 2022). We did however find α-synuclein associated with 5HTT positive synapses and, as with dopaminergic synapses, a positive correlation between SNAP-25 volume and α-synuclein volume. This change in 5HT synapses may indicate dysfunction of 5HT terminals without a change in serotonergic innervation in the sgACC since we saw no change in fibre density or 5HTT protein levels. This correlates with other studies that showed no association with 5HTT levels with depression or anxiety in PD (Qamhawi et al. 2015). Future studies should however determine the effects of α-synuclein pathology in DLB in the brainstem and midbrain serotonergic system using appropriate stereological approaches and anatomical markers.

There has been considerable use of serotonin transporter ligands in PD and to a lesser extent DLB. In drug naïve PD patients, there were no changes in non-apathetic patients but reductions in [(11)C-N,N-dimethyl-2-(-2-amino-4-cyanophenylthio)-benzylamine (DASB)] retention in apathetic patients in the sgACC and supragenual cingulate, medial orbitofrontal cortex, anterior caudate and ventral striatum when compared to non-apathetic PD or controls suggesting reduced serotonergic innervation (Maillet et al. 2016). Similarly, reduction of SERT assessed with DASB in caudate and ACC is seen in early stage PD and throughout the disease course (Politis, Wu, Loane, Turkheimer, et al. 2010; Albin et al. 2008; Guttman et al. 2007; Maillet et al. 2021; Wilson et al. 2018; Politis, Wu, Loane, Kiferle, et al. 2010) and at post mortem (Kish et al. 2008). In contrast, in one small study of depressed early stage PD patients, increase in DASB retention was observed in several cortical areas although there is generally no change in DASB retention in ACC in MDD (Boileau et al. 2008; Kambeitz and Howes 2015). Studies in DLB using in vivo imaging have been restricted to the use of the combined DAT and SERT ligand ^123^I-N-ω-fluoropropyl-2β-carbomethoxy-3β-(4-iodophenyl) nortropane (^123^I-FP-CIT) where the assumption is that cortical retention only represents 5HT innervation despite the presence of DA terminals. Studies with ^123^I-FP-CIT have shown either reduced cortical SERT (Nicastro et al. 2020; van der Zande et al. 2020; Pilotto et al. 2019), regional reductions but not within the cingulate (Takahashi et al. 2021), or no cortical changes (Joling et al. 2019). No reduction in SERT was seen in the ACC at post mortem in advanced PD with dementia, although other brain regions such as caudate showed reductions (Buddhala et al. 2015). Similarly, in DLB compared to control or non-depressed patients, SERT assessed with cyanoimipramine showed preserved cortical binding in depressed DLB donors although an overall reduction is suggested (Ballard et al. 2002; Francis 2009). Whilst these previous studies have shown variable outcomes, based on the current findings variable, SERT reductions in DLB may be mild or absent and contributions to development of depression are potentially limited.

Decreased serotonin 5HT1_A_ binding has been shown in the ACC in MDD patients using PET (Wang et al. 2016) however, we observed no alteration of 5HT1_A_ protein levels in DLB. This contrasts with findings using the 5HT1a receptor ligand 8-Hydroxy-2-Dipropylaminotetralin that showed increased binding but with a change in K_D_ in both DLB and PD (Sharp et al. 2008). This may suggest that whilst there is a change in receptor occupancy, this is due to changes in protein conformation but not protein levels as seen in the current study. However, our finding of a decrease in 5HT2_A_ protein in sgACC in DLB donors with depression corresponds with reduced 5HT2_A_ binding in ACC and DLPFC in treatment resistant MDD (Baeken, De Raedt, and Bossuyt 2012) and reduced [^3^H]-ketanserin binding in DLB and PDD (Cheng et al. 1991; Perry et al. 1990). These changes in 5HT2_A_ but not 5HT1_A_ may suggest selective and region specific serotonergic receptor changes that may also benefit from selective targeting. The use of Pimavanserin, a 5HT2a selective inverse agonist and antagonist, has been explored in PD for the treatment of psychosis including delusions and hallucinations caused by dopaminergic medication (Meltzer et al. 2010; Cummings et al. 2014). Despite mixed benefits in MDD (Dirks et al. 2022; Fava et al. 2019), a single small open label study of Pimavanserin for treatment of depression in PD in the absence of psychosis showed delayed benefit in an 8-week trial with 60% of patients showing improved mood (DeKarske et al. 2020). Longer-term use of Pimavanserin may though show reduced efficacy (Ahmed et al. 2022). The possibility of using Pimavanserin in the treatment of depression in may therefore be warranted since Pimavanserin shows some effect in reducing psychosis in certain DLB patients (Horn et al. 2019).

In summary, these results suggest that there is primarily a reduced dopaminergic drive associated with depression in DLB, with reduced dopaminergic innervation in the sgACC, along with reduced levels of dopaminergic markers and receptors, and changes in dopaminergic synaptic function. Careful treatment with selective dopaminergic agonists or positive allosteric modulators may therefore be of benefit in alleviating depressive symptoms in DLB.

