## Supplementary Material for "Dopaminergic Changes in the Subgenual Cingulate Cortex in Dementia with Lewy Bodies Associates with Presence of Depression"

### Supplementary Table 1. Antibodies used in the study.

| Antigen | Host/Ig | Dilution | Source | Product Code |
|---|---|---|---|---|
| 5HT1A | pRb (IgG) | 1:1000 (DB) | Abcam | ab227165 |
| 5HT2A | pRb (IgG) | 1:100 (DB) | Abcam | ab66049 |
| 5HT3B | pRb (IgG) | 1:500 (DB) | Abcam | ab39629 |
| 5HTT | mMs (IgG1) | 1:1000 (DB), 1:500(STED) | MAb Technologies | ST51-2 |
| Tryptophan hydroxylase 2 (TPH-2) | mMs (IgG1) | 1:500 (DB) | Atlas Antibodies | AMAb91108 |
| Dopamine transporter (DAT) | mMs (IgG1) | 1:250 (DB), 1:20 (STED) | Atlas Antibodies | sc32258 |
| Tyrosine Hydroxylase (TH) | pRb (IgG) | 1:500 (DB) | Abcam | ab112 |
| D2DR | pRb (IgG) | 1:500 (DB) | Abcam | ab130295 |
| D3DR | mRb (IgG) | 1:1000 (DB) | Abcam | ab155098 |
| D4DR | mMs (IgG1) | 1:500 (DB) | Santa Cruz Biotechnology | sc136169 |
| DOPA Decarboxylase (DDC) | mMs (IgG1) | 1:500 (DB) | Atlas Antibodies | AMAb91089 |
| SNAP25 | pSh (IgG) | 1:500 (STED) | LifeSpan BioSciences | LS-C94834 |
| α-synuclein (KM51) | mMs l (IgG1) | 1:50 (IHC) | Leica Biosystems | ASYN-L |
| α-synuclein (5G4) | mMs (IgG1) | 1:4500 (IHC) | Analytik Jena | 847-0102004001 |
| α-synuclein (s129) | pRb (IgG) | 1:500 (DB), 1:500 (F-IHC) | Abcam | ab168381 |
| Amyloid-beta (4G8) | mMs ( IgG2b) | 1:12000 (IHC) | BioLegend | 800701 |
| Tau (AT8) | mMs (IgG1) | 1:8000 (IHC) | Thermo Fisher | MN1020 |
| HuD (neuronal marker) | mMs (IgG2a) | 1:1000 (IHC) | Santa Cruz Biotechnology | sc48421 |
| Donkey Anti-Sheep, 594 conjugate | Sheep (IgG) | 1:50 (STED) | Life Technologies | A11016 |
| Goat Anti-Rabbit, Atto 647N conjugate | Rabbit (IgG) | 1:50 (STED) | Sigma-Aldrich | 40839 |
| Goat Anti-Mouse, 532 conjugate | Mouse (IgG) | 1:50 (STED) | Life Technologies | A-11002 |

DB – dot blot; IHC – immunohistochemistry; m – monoclonal; p – polyclonal; Ms – mouse; Rb – rabbit, Gt – goat, Dk – donkey.

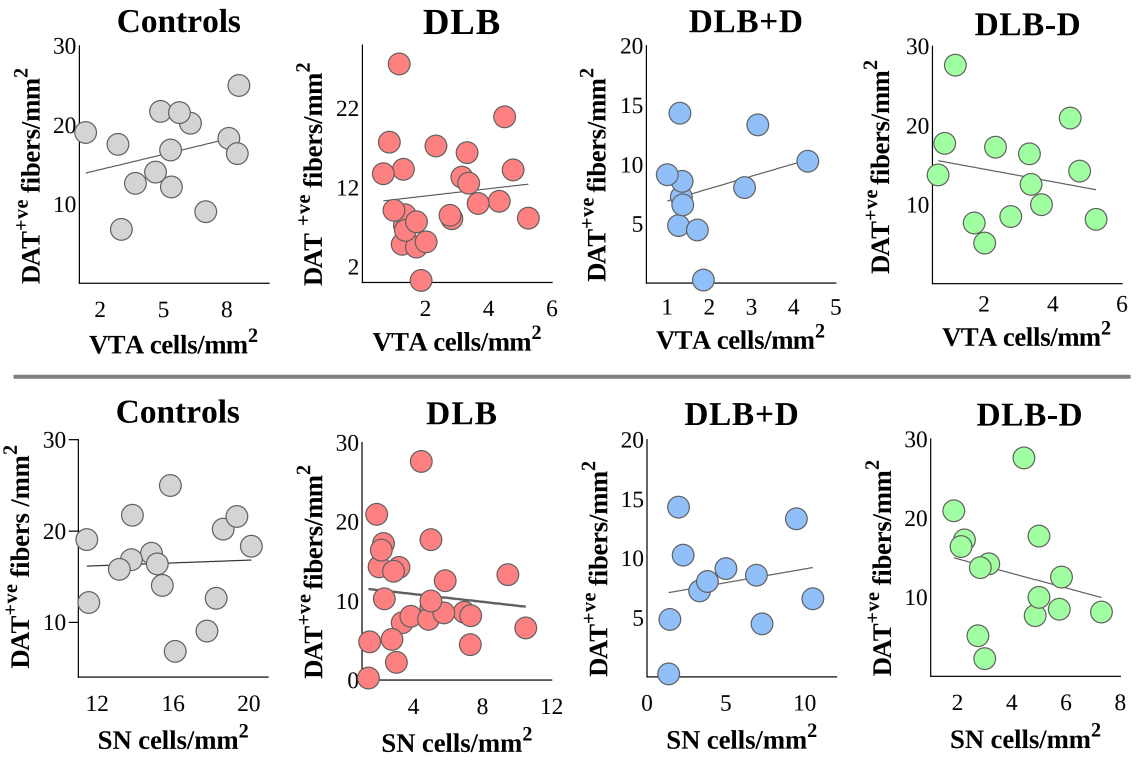


**Supplementary Figure 1. Effect of dopaminergic neurons in VTA and SN on dopaminergic fibers in sgACC.**

**Top)** Correlations between DAT+ve fibre density in sgACC and dopaminergic neurons in VTA; **Bottom)** and dopaminergic neurons in SN within controls, DLB cases with and without depression.

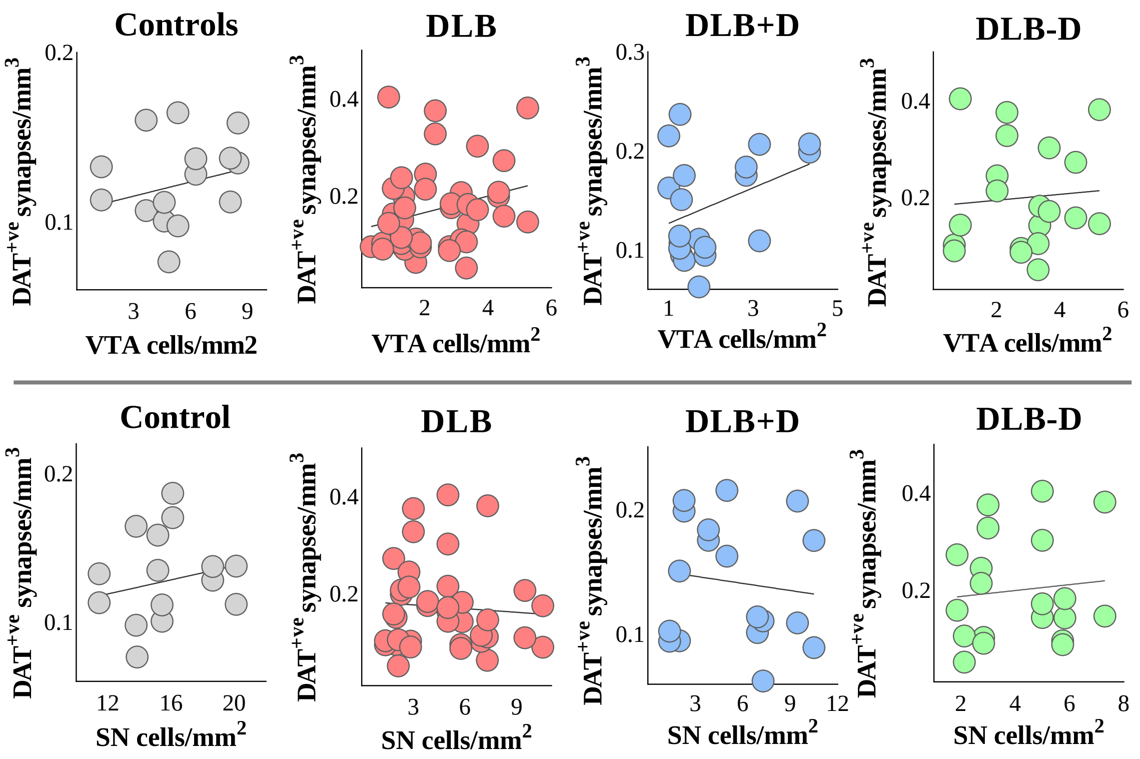


**Supplementary Figure 2. Effect of dopaminergic neurons in VTA and SN on dopaminergic synapses in sgACC.**

**Top)** Correlations between DAT+ve synapses in sgACC and dopaminergic neurons in VTA; **Bottom)** and dopaminergic neurons in SN within controls, DLB cases with and without depression. Significant positive correlation between dopaminergic neurones in VTA and DAT positive synapses in sgACC in DLB cases with depression (*$p<0.05$).

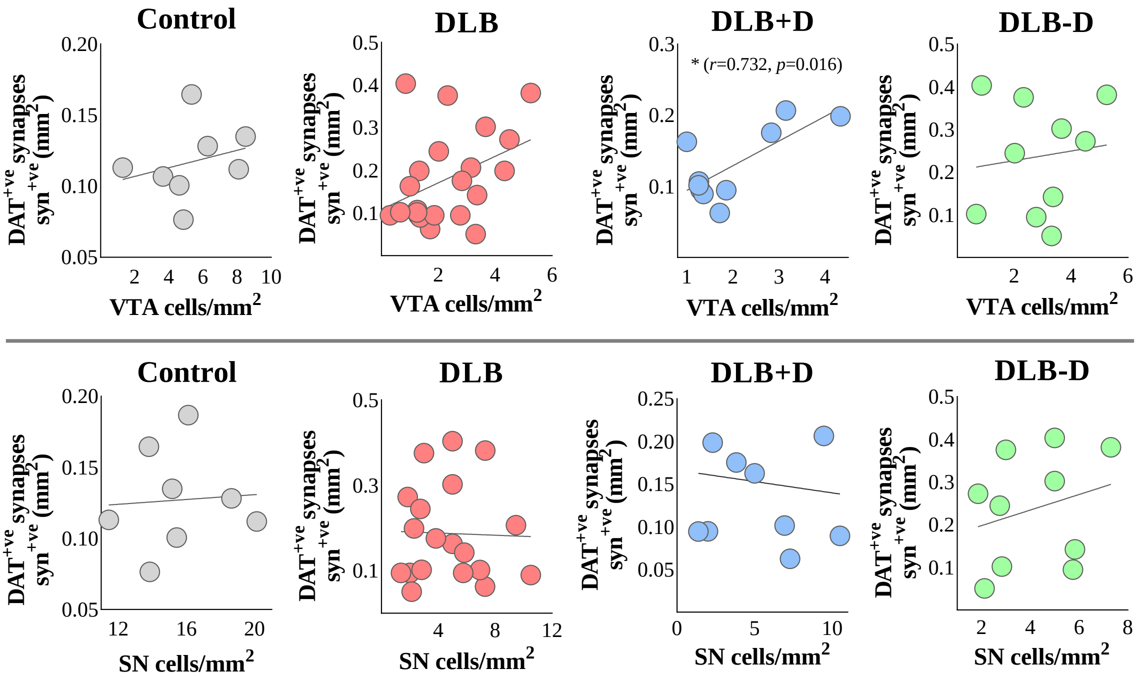


**Supplementary Figure 3. Effect of dopaminergic neurons in VTA and SN on α-synuclein positive dopaminergic synapses in sgACC.**

**Top)** Correlations between α-synuclein positive DAT+ve synapses in sgACC and dopaminergic neurons in VTA; **Bottom)** and dopaminergic neurons in SN within controls, DLB cases with and without depression. Significant positive correlation between dopaminergic neurones in VTA and α-synuclein positive DAT positive synapses in sgACC was observed in DLB cases with depression (*$p<0.05$).
